# Novel Imaging Tools to Study Mitochondrial Dynamics in *Caenorhabditis elegans*

**DOI:** 10.1101/2024.07.16.603730

**Authors:** Miriam Valera-Alberni, Pallas Yao, Silvia Romero-Sanz, Anne Lanjuin, William B. Mair

**Affiliations:** Dept. Molecular Metabolism, Harvard TH Chan School of Public Health, Massachusetts, USA

## Abstract

Mitochondria exhibit a close interplay between their structure and function. Understanding this intricate relationship requires advanced imaging techniques that can capture the dynamic nature of mitochondria and their impact on cellular processes. However, much of the work on mitochondrial dynamics has been done in single celled organisms or in vitro cell culture. Here, we introduce novel genetic tools for live imaging of mitochondrial networks in the nematode *Caenorhabditis elegans*, addressing a pressing need for advanced techniques in studying organelle dynamics within live intact multicellular organisms. Through a comprehensive analysis, we directly compare our tools with existing methods, demonstrating their advantages for visualizing mitochondrial morphology and contrasting their impact on organismal physiology. We reveal limitations of conventional techniques, while showcasing the utility and versatility of our approaches, including endogenous CRISPR tags and ectopic labeling. By providing a guide for selecting the most suitable tools based on experimental goals, our work advances mitochondrial research in *C. elegans* and enhances the strategic integration of diverse imaging modalities for a holistic understanding of organelle dynamics in living organisms.

## Introduction

Mitochondria, essential organelles in eukaryotic cells, exhibit a dynamic and interconnected network architecture crucial for their functional adaptability. This intricate network is governed by coordinated processes such as membrane fusion, fission, and mitochondrial trafficking, collectively referred to as mitochondrial dynamics. The ability of mitochondria to undergo morphological changes is pivotal for their functional capacity to respond to nutrient availability, molecular signals, and propagate mitochondrial genomes (Liesa & Shirihai, 2013). Disruptions in this network organization have been linked to mitochondrial dysfunction, thereby contributing to the onset of various pathophysiological conditions, including neurodegenerative diseases, cardiovascular disorders, cancer, or metabolic diseases (Chen *et al*, 2023). Consequently, research into mitochondrial dynamics have become integral to comprehending mitochondrial function in disease states. Despite this growing awareness, a significant gap remains in the availability of optimal imaging tools to visualize mitochondrial morphology in multicellular living organisms. Much of the work to date has been performed in single-celled organisms such as *Saccharomyces cerevisiae* or using in vitro cell culture. Although these systems have many advantages for understanding the molecular mechanisms governing mitochondrial regulation, they are more limited in their utility when it comes to physiology. Conversely, where mouse models support examination of how changes to mitochondrial dynamics impact physiology and pathophysiology, in vivo imaging is a challenge. This study aims to fill this gap by introducing novel imaging tools tailored for the nuanced visualization of mitochondria in the nematode worm *Caenorhabditis elegans*, offering new opportunities for the study of mitochondrial dynamics in disease conditions and longitudinally throughout an animal’s life.

Powerful genetic approaches have expedited the molecular characterization of multiple mitochondrial processes, making *C. elegans* an ideal animal model for understanding the organismal roles of mitochondria (Onraet & Zuryn, 2024). Many fundamental genetic, biochemical, and structural features of mitochondria remain conserved between mammals and *C. elegans* (Campbell & Zuryn, 2024; Ha *et al*, 2022), including key components of the mitochondrial dynamics machinery and receptor complexes of the outer (OMM) and inner (IMM) mitochondrial membranes. In line with this, *C. elegans* provides a particular useful platform for visualizing mitochondrial morphology in vivo. As compared to single cell models such as yeast and cell culture, *C. elegans* offers a complex organization of differentiated tissues, including a defined nervous system, hypodermis, gastrointestinal system, muscle, and germ lineage in which to study cell-type specific and cell non-autonomous features of mitochondrial function (Burkewitz *et al*, 2015; Onraet & Zuryn, 2024). Further, the transparency of its cuticle allows for non-invasive, high-resolution imaging of mitochondria in live animals. This is in contrast with studies in mice, where changes in mitochondrial morphology are derived from biopsied tissues extracted from sacrificed animals. Imaging in *C. elegans* is conducted without tissue fixation, thus allowing the visualization of real-time mitochondrial dynamics under distinct physiological conditions, which facilitates longitudinal studies. Indeed, the short lifespan of *C. elegans* enables researchers to conduct aging experiments that otherwise would require substantially more time in mice (mean lifespan of 2 years) (Apfeld & Alper, 2018). Finally, the genetic tractability of worms allows targeted manipulation of mitochondrial components, enabling precise investigations into the molecular mechanisms governing mitochondrial dynamics.

The potential of *C. elegans* for cell biology in vivo, however, has been hindered by a lack of development of tools to visualize organelles for quantitative microscopy. We aimed to address this gap by engineering a suite of novel transgenic stains to study mitochondrial dynamics in a living animal. With the advent of CRISPR knock in gene editing, sophisticated approaches to generate new tools and visualize endogenous proteins are now feasible. We demonstrate that these new tools overcome some of the caveats posed by previously used methods to visualize mitochondria in *C. elegans*. Ultimately, we provide a comparison of past and current tools, highlighting specific caveats and giving researchers a data led approach to determine best imaging practices. Together, we believe that the strains presented here provide the foundation for in vivo cell biology studies of mitochondria in *C. elegans*.

## Results

### Limitations of current tools to study mitochondrial structure in *C. elegans*

Increasing attention has been directed towards exploring the vital role of mitochondrial dynamics in influencing mitochondrial function and, consequently, its impact on organismal physiology. Advanced imaging tools are essential for the analysis of mitochondrial dynamics, yet the instruments employed to investigate mitochondrial morphology in *C. elegans* remain outdated. The primary indicator of mitochondrial morphology in most publications involving *C. elegans* was established in 2006, a mitochondrial-targeted green fluorescent protein (GFP^mt^) under the control of the muscle-specific myo-3p promoter (Benedetti *et al*, 2006). However, this tool presents several caveats that hinder its utility in cell biology studies. First, this model was generated using extrachromosomal plasmid arrays and integrated in the genome via gamma irradiation in unknown copy number and at a random locus (Benedetti *et al*, 2006). Variability in baseline expression levels make quantitative microscopy nearly impossible within the same strain, and especially across multiple conditions. In addition, possibly due to off-target effects of gamma irradiation or the high expression levels of the multicopy array, the mito-GFP strain displays slight developmental delay and reduced brood size (**Fig 1A**). Second, as a mito-targeted fluorophore that is non-integrated into the membrane or constitutively residing in the lumen, visualization of mitochondrial networks relies on the fidelity of mitochondrial protein import. Mitochondrial protein import of GFP^mt^ is inconsistent even in young day 1 worms, as shown by inadequate GFP labeling of the mitochondrial network and cytosolic signal in some animals (**Fig 1B**). In addition, lifespan analysis reveals that the mito-GFP strain is long lived (**Fig 1C**) suggesting broader effects on physiology and mitochondrial function. Despite the fact that extrachromosomal arrays offer a reliable and fast means to express transgenes in *C. elegans*, they also have limitations. Since extrachromosomal arrays are semistable, only a fraction of the animals in a transgenic extrachromosomal array line express and propagate the transgene (Praitis *et al*, 2001). This leads to the laborious task of selecting the animals with the integrated transgene before starting any experimental approach. In addition, variations in array copy number and repeat sequence silencing can lead to heterogeneity in expression level and pattern of the transgene between individuals. Many existing tools in *C. elegans*, including those made by us previously, use extrachromosomal TOMM-20^1-49^::GFP arrays that selectively visualize mitochondria within muscle, facilitated by a muscle-specific promoter (**Fig EV1A**). However, this does not permit imaging of mitochondria in other tissues (**Fig EV1A**). In addition to the labor-intensive process of individually selecting each GFP-positive worm, these animals exhibit minor developmental delay and reduced brood size (**Fig EV1B**), as well as slight extension in lifespan (**Fig EV1C**).

**Figure 1.**
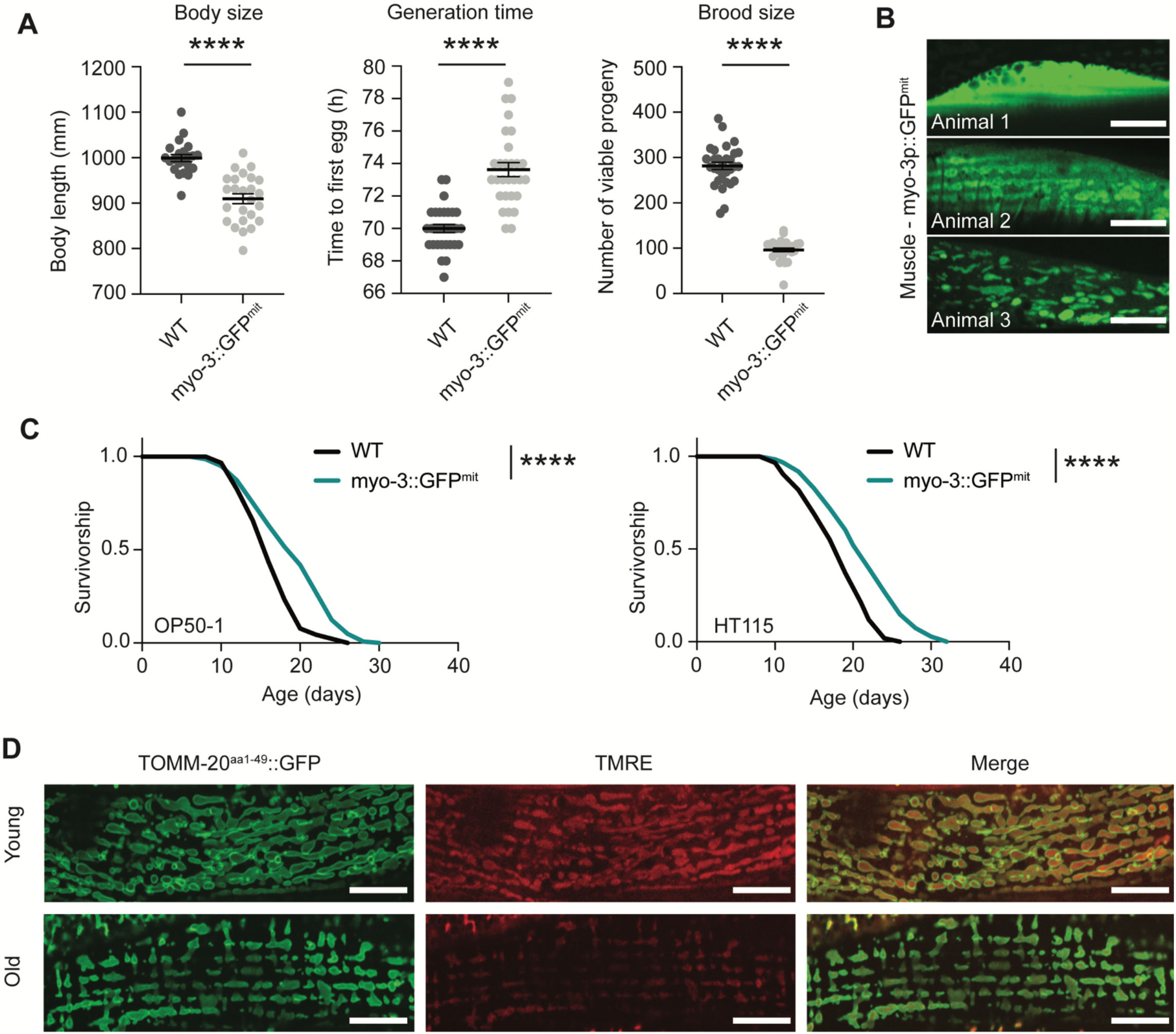
Limitations of current mitochondrial imaging tools in *C. elegans*. A. Healthspan analysis of the myo-3::GFP^mit^ strain compared to wild-type N2 worms, showing body size (left graph), generation time (middle), and brood size (right). Body size measurements were performed with 20-25 individual worms. Generation time and brood size measurements were performed with 30 worms pooled from two independent experimental repeats. All values are presented as mean ± SEM. ****p < 0.0001 versus the respective WT group (two-tailed Student’s t test) B. Representative fluorescence pictures depicting mitochondrial networks in the body wall muscle of 3 different worms on day 1 of adulthood. Same laser exposure (200 ms) was used for the 3 images. The upper panel displays a muscle cell with a predominantly cytosolic GFP signal from myo-3p::GFP^mit^, while the middle and bottom panels show a gradual labeling of mitochondria by GFP. Scale bar: 10 μm. C. Survival curve of the myo-3::GFP^mit^ strain compared with WT in OP50-1 (left lifespan) and HT115 (right lifespan) bacteria. ****p < 0.0001 indicating significant difference compared to WT animals by one-way ANOVA with Tukey’s multiple comparisons test. D. Worms expressing TOMM-20^aa1-49^::GFP were treated with TMRE and examined on the second day of adulthood. Subsequently, both green (GFP) and red (TMRE) signals were captured in both young (day 1) and aged (day 8) worms. Scale bar: 10 μm.

Inconsistency in mitochondrial labelling is also true for mitochondrial dyes, such as tetramethylrhodamine ethyl ester (TMRE), which are impacted by mitochondrial membrane potential (Gökerküçük *et al*, 2020; Chazotte, 2009). Depolarized mitochondria, which tend to appear in disease-prone conditions such as aging (Berry *et al*, 2023), display decreased membrane potential and fail to sequester TMRE. Therefore, staining with TMRE to contrast mitochondrial network morphology across conditions can be confounded by technical variability (**Fig 1D**). Moreover, staining by TMRE varies across tissues due to its infiltration, proving highly effective in the hypodermis or the intestine but showing lower infiltration in muscle and head tissues (**Fig 2**). Further, off-target effects such as dye aggregation or quenching in the mitochondria can interfere with the experimental approach.

**Figure 2.**
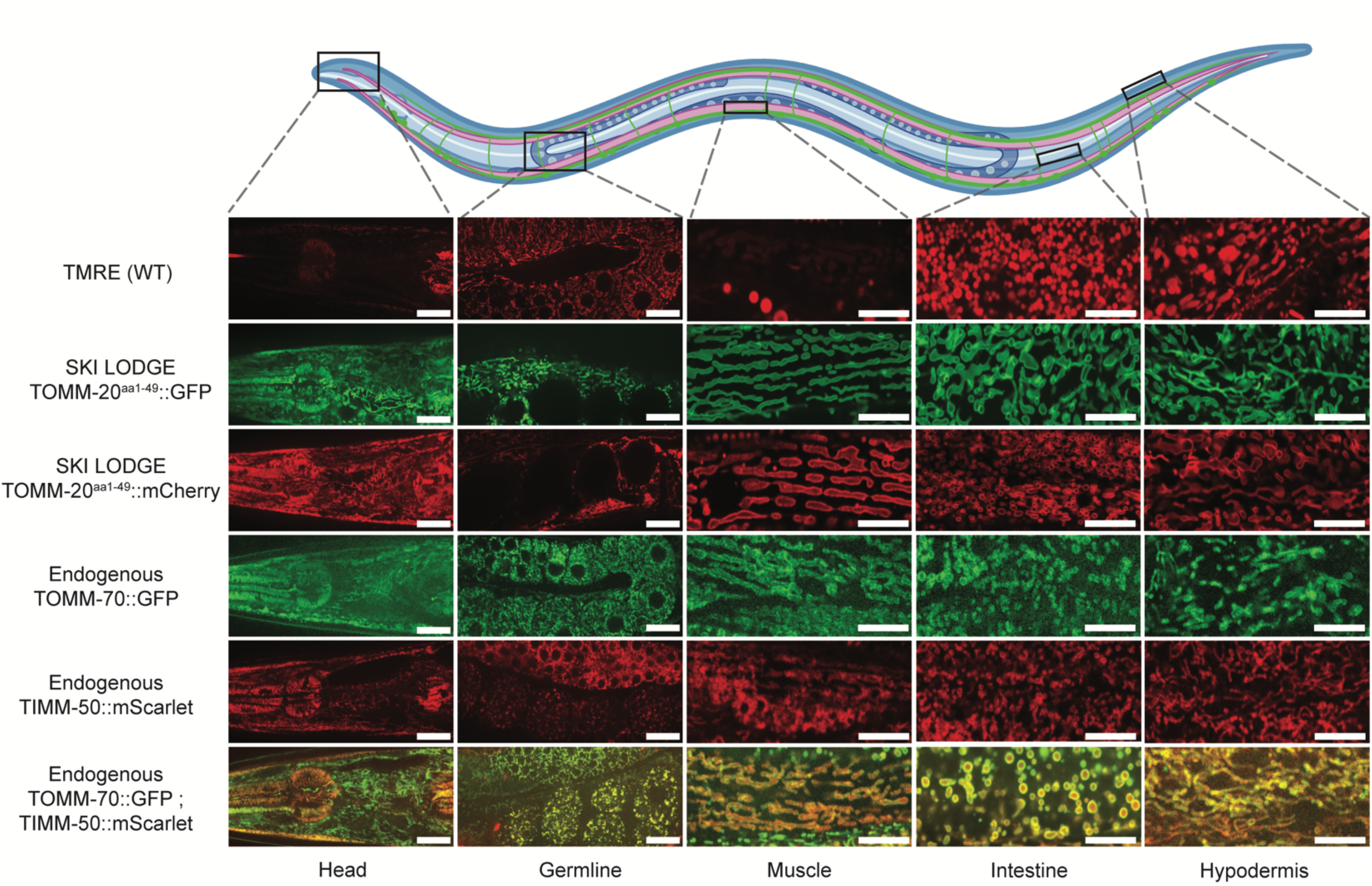
Improved mitochondrial reporter tools in *C. elegans*. Comparative analysis of mitochondrial visualization techniques in various tissues of *C. elegans*. The upper panel of images demonstrates the variable staining efficacy of TMRE in wild-type N2 worms. The novel SKI LODGE strains, expressing the TOMM-20 transmembrane domain fragment fused to GFP or mCherry, respectively, under the *eft-3p* promoter, exhibit more uniform labeling. Endogenous mitochondrial strains include, GFP expression on the translocase TOMM-70, or mScarlet tagging on the translocase of the inner mitochondrial membrane TIMM-50, respectively. Dual labeling of both mitochondrial membranes is achieved in a strain expressing GFP on the OMM and mScarlet on the IMM. Scale bar (head, germline): 500 μm. Scale bar (muscle, intestine, hypodermis): 300 μm.

To circumnavigate these caveats, we generated several new fluorescence reporters that are expressed somatically or ubiquitously to visualize mitochondrial morphology in live *C. elegans* via two strategies: first, via CRISPR-assisted single copy knock in of somatically expressed TOMM-20^1-49^::GFP/mCherry and second, via CRISPR in frame integration of fluorophores into the loci of genes encoding endogenous mitochondrial membrane proteins.

### Visualizing mitochondria across multiple tissues in *C. elegans*

Advancements in precision gene editing techniques, particularly in the context of manipulating genes within *C. elegans*, have yielded improved tools to probe mitochondria. We generated novel strains by tagging components of the outer mitochondrial membrane (OMM) to fluorescence markers such as GFP, mCherry or mScarlet. First, we generated two novel strains carrying the 1-49aa transmembrane domain fragment of the OMM translocase TOMM-20 (TOMM-20) fused to GFP or mCherry, driven by the *eft-3* promoter for ubiquitous tissue expression (TOMM-20^1-49^::GFP/mCherry). Using CRISPR/Cas9 editing via the SKI LODGE method (Silva-García *et al*, 2019), this construct was integrated in single copy at a defined intergenic region on Chromosome V, previously characterized to give stable expression with no silencing (Silva-García *et al*, 2019). Compared to the mito-GFP strain or staining with dyes, these new lines show more consistent expression and accurate labeling of mitochondrial networks in worms. In these strains, the visualization of mitochondria reaches high intensity and resolution across various tissues including muscle, intestine, and hypodermis (**Fig 2**), and the neuronal network in the worm’s head. Of note, the ectopic *eft-3* promoter in the SKI LODGE system is not expressed in the germline. In addition, the TOMM-20^1-49^::GFP/mCherry markers in the SKI LODGE strains do not give any insight into the relative abundance of endogenous mitochondria proteins in the worm. To circumvent these issues, we also used CRISPR to generate a strain with GFP fused to the C-terminus of the endogenous OMM translocase TOMM-70 or, alternatively, mCherry tagged to the translocase of the endogenous inner mitochondrial membrane (IMM), TIMM-50 (Paix *et al*, 2015). Finally, we crossed these strains together to be able to visualize the OMM and IMM together in the same animal, which proves notably useful for studies requiring differentiation between processes exclusive to each of these mitochondrial membranes (**Fig 2**). Having generated these new tools, we then sought to further characterize their utility and effect on animal physiology.

### Single Copy Ectopic OMM Markers

Using ectopic promoters to drive OMM markers has several advantages. Endogenous mitochondrial membrane proteins remain wild-type (WT), while the higher expression of promoters such as *eft-3* even at single copy increases signal for microscopy. Indeed, a direct comparison of fluorescence intensity between SKI LODGE strains (TOMM-20^1-49^::GFP/mCherry) and endogenous strains (TOMM-70::GFP and TIMM-50::mScarlet, respectively) revealed higher intensity levels in the former (**Fig 3A**). This heightened fluorescence intensity facilitates confocal imaging, providing improved clarity and ease of analysis. Tagging fluorophores to mitochondrial membranes, membrane proteins or proteins involved in mitochondrial remodeling can affect mitochondrial function (Montecinos-Franjola *et al*, 2020). Reduced mitochondrial function reduces body size, delays development, and decreases reproduction in *C. elegans* (Durieux *et al*, 2011; Dillin *et al*, 2002). To determine if the new SKI LODGE ubiquitous TOMM-20^1-49^ reporter causes phenotypic differences due to the fluorophore or off-target toxicity, we therefore measured several healthspan parameters, such as growth, development, and fertility. While the TOMM-20^1-49^::GFP animals exhibited consistency in healthspan parameters, including body size, generation time, and brood size, as compared to the wild-type N2 strain, TOMM-20^1-49^::mCherry worms displayed slightly increased generation time and reduced brood size (**Fig 3B**). This suggests that the introduction of the GFP/mCherry tag to TOMM-20 does not adversely impact these crucial health-related metrics. Lifespan analysis on *E. coli* OP50-1 indicated no significant changes compared to wild-type animals, although a slight reduction in lifespan was observed on HT115 (**Fig 3C**). It’s important to note that NG plates containing antibiotics were utilized when using HT115 bacteria. Overexpression of reporters through the SKI LODGE system may therefore exhibit sensitivity to antibiotics, potentially influencing lifespan outcomes.

**Figure 3.**
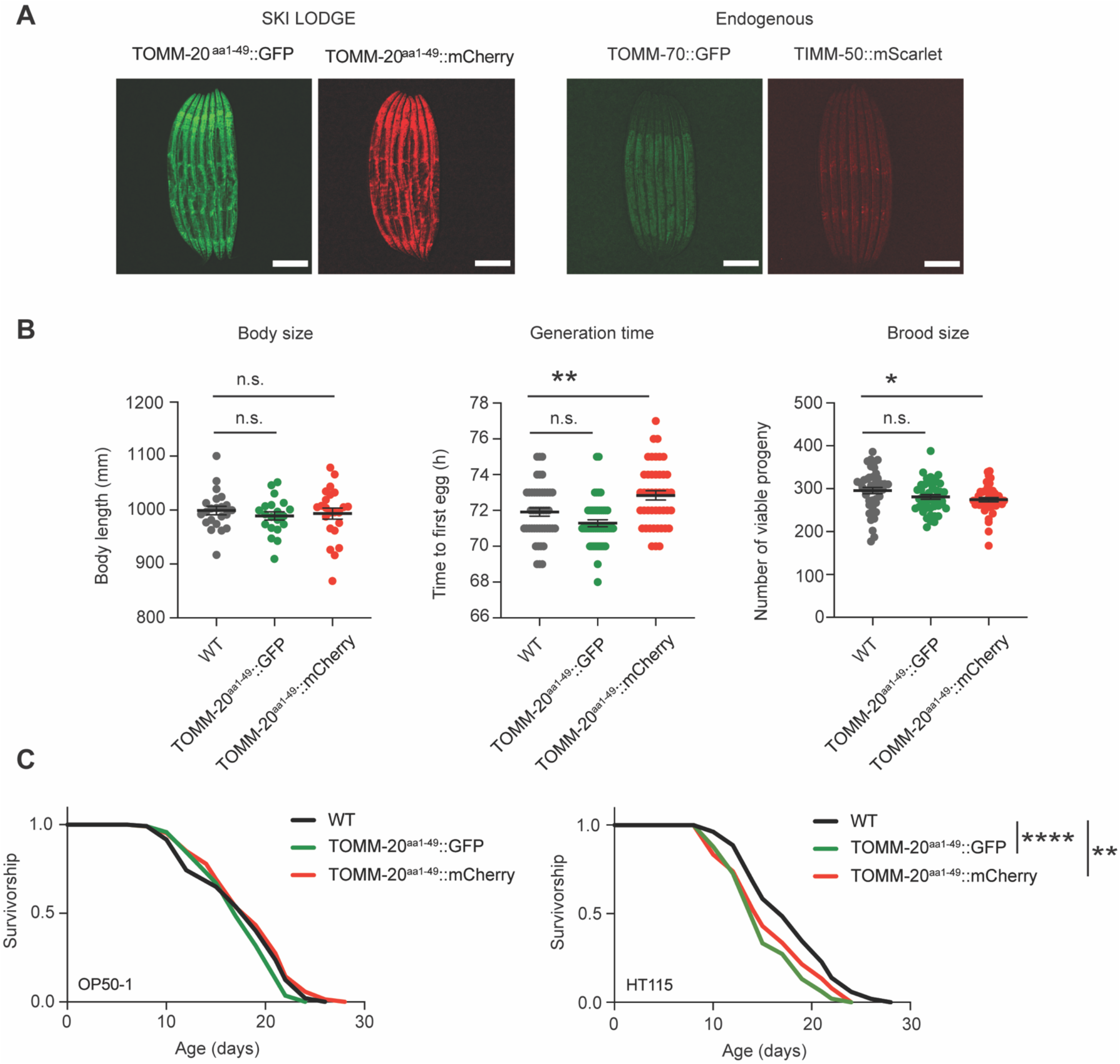
Novel TOMM-20 reporter strains generated using SKI LODGE in *C. elegans*. A. Fluorescence image of the SKI LODGE TOMM-20^aa1-49^::GFP and TOMM-20^aa1-49^::mCherry worms, respectively, as well as the TOMM-70::GFP and TIMM-50::mScarlet worms at day 1 of adulthood. Laser exposure: 65 ms. Scale bar: 200 μm. B. Healthspan analysis of SKI LODGE strains compared to wild-type N2, showing body size (left), generation time (middle), and brood size (right). Body size measurements were performed with 25 individual worms. Generation time and brood size measurements were performed with 45 worms pooled from three independent experimental repeats. All values are presented as mean ± SEM. One-way ANOVA followed by a multiple comparisons test with the mean of the control (WT) group was performed to identify statistically significant data. *p < 0.05, **p < 0.01. C. Survival of SKI LODGE strains compared with WT in OP50-1 (left lifespan) and HT115 (right lifespan) bacteria. **p < 0.01 indicating significant difference between WT animals and TOMM-20^aa1-49^::mCherry worms, ***p < 0.001 represents significant difference between WT and TOMM-20^aa1-49^::GFP worms by one-way ANOVA with Tukey’s multiple comparisons test. See Table EV3 for detailed lifespan statistics.

### Fluorescent tagging of endogenous mitochondrial membrane proteins

Since expression of TOMM-20^1-49^::GFP or mCherry in the SKI LODGE strains is driven by an ectopic promoter, it cannot be used to determine relative abundance of endogenous mitochondria proteins in the worm. Having used CRISPR to generate a strain expressing GFP on the C-terminus of endogenous outer membrane translocase TOMM-70 (Paix *et al*, 2015), we sought further characterize it. This strain facilitates imaging of mitochondrial networks across all tissues, including the germline, which was not possible in previous extrachromosomal strains or the SKI LODGE strain. In line with this, TOMM-70 displays high expression in the germline of *C. elegans*, as compared to other tissues such as the muscle, intestine, or the hypodermis (**Fig 4A**). The TOMM-70::GFP strain did not exhibit any alterations in body size, generation time or brood size as compared to WT N2 worms (**Fig 4B**). Similarly, the median lifespan of the TOMM-70::GFP strain was comparable to that of the WT animals. Overall, these observations suggest that the endogenous GFP tag on TOMM-70 does not lead to any overt physiological abnormalities. Contrarily, physiological effects were observed in the TIMM-50::mScarlet animals including altered growth, slowed development and decreased reproductive capacity (**Fig 4B**). Despite this, the TIMM-50::mScarlet animals had no difference in median lifespan relative to WT animals fed on OP50-1 or HT115 bacteria. This was also the case for the double TOMM-70::GFP ; TIMM-50::mScarlet animals, which showed increased generation time and reduced brood size (**Fig 4B and 4C**).

**Figure 4.**
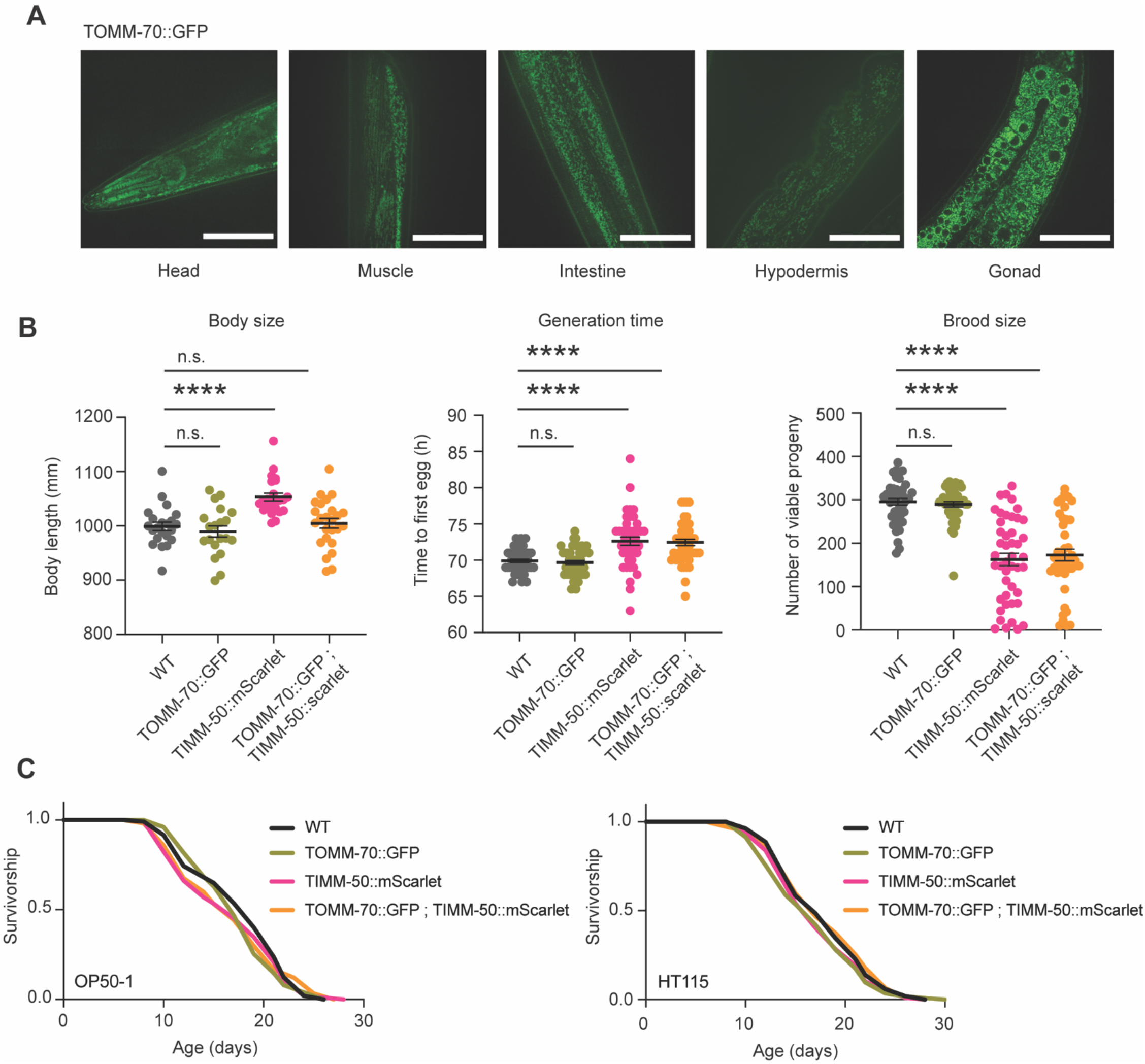
Endogenous tagging of mitochondrial membrane proteins in *C. elegans*. A. Visualization of relative endogenous expression levels of TOMM-70 through the intensity of GFP in various *C. elegans* tissues of the TOMM-70 endogenous strain. Scale bar: 50 μm. B. Comparative healthspan analysis among TOMM-70::GFP, TIMM-50::mScarlet, and the double TOMM-70::GFP; TIMM-50::mScarlet strains, respectively, in relation to wild-type N2 worms. Parameters such as body size (left graph), generation time (middle), and brood size (right) are presented for these strains. Body size measurements were performed with 25 individual worms. Generation time and brood size measurements were performed with 45 worms pooled from three independent experimental repeats. All values are presented as mean ± SEM. ****p < 0.0001 (One-way ANOVA followed by multiple comparisons test) between the indicated groups. D. Survival of endogenous strains compared with WT in OP50-1 (left lifespan) and HT115 (right lifespan) bacteria.

In summary, CRISPR/Cas9 facilitated the generation of new tools to better study mitochondrial networks and dynamics via fluorescence tagging of mitochondrial outer and inner membrane components, notably with the TOMM-70::GFP strain providing a nuanced view of mitochondrial expression levels across tissues. These tools hold promise for researchers studying mitochondrial morphology changes in live *C. elegans*.

## Discussion

*C. elegans* represent a unique middle ground between single cell systems and mammals to study organelle biology (Onraet & Zuryn, 2024). No one tool is best equipped to take on this challenge, and instead researchers must decide which tool best serves them depending upon their scientific question, resources, and timeline (**Fig EV6**). Several methods exist that do not require gene editing, including electron microscopy, immune-fluorescence, and cellular dyes such as MitoTracker and TMRE. However, although these all have unique advantages, they all require fixing or termination of the sample, which limits their utility for longitudinal and noninvasive live imaging. The introduction of novel *C. elegans* strains with fluorophore-tagged components of the outer and inner mitochondrial membrane marks a significant advancement in our ability to investigate mitochondrial dynamics within intact living organisms. Here, we introduce novel fluorophore-based genetic tools for imaging mitochondrial networks in *C. elegans*, addressing the limitations of existing tools (**Table EV1**).

Using targeted gene-editing strategies to generate fluorescent reporters of mitochondrial membranes offers substantial advantages over extrachromosomal methods as it allows stable and uniform expression within an isogenic population of animals. This is critical for accurate quantitative analysis of mitochondrial dynamics. It also improves upon gamma irradiation methods of integrating edits into the *C. elegans* genome as it has less off-target effects, and bypasses the need for co-selection. Our SKI LODGE strains have increased fluorophore intensity, which is homogeneous across different tissues and allows straightforward imaging using basic microscopes without much need for imaging processing. For analysis of endogenous protein expression levels, strains such as the TOMM-70::GFP or TIMM-50::mScarlet can be employed, albeit with reduced fluorophore intensity and the need for fine-tuning of higher resolution image acquisition.

How the tool impacts the function of mitochondria themselves or organismal physiology is also a critical consideration, especially when conducting longevity studies and imaging late in life. Whereas the mito-GFP line showed significant reduction in body size, brood size, and delayed development, neither the SKI LODGE lines show overt alterations in healthspan indicators. Although the mean generation time of the SKI LODGE TOMM-20^1-49^::mCherry strain (70.84 hrs ± 0.3266) was significantly different as compared to WT (69.91 hrs ± 0.3474) animals, this effect is negligible. Similarly, this line shows mildly reduced brood size. Moreover, both SKI LODGE strains display a slight lifespan reduction when aged on NG plates with antibiotics fed on HT115. Maintenance of *C. elegans* in the laboratory standardly includes the removal of spontaneously occurring microbial contaminants to limit experimental variations, since diet, including the presence of contaminants, has been shown to exert potent influences over animal physiology, such as development, reproduction, healthspan, and longevity (Stuhr & Curran, 2020). This is routinely achieved by use of a single or combination of antibiotics. Bacterial food sources have also been engineered to carry specific resistance for selection, and this is particularly relevant for RNA interference studies in *C. elegans*, where worms are fed RNAi-containing bacteria. In these studies, carbenicillin is consistently included to guarantee the exclusive presence of RNAi or control bacteria carrying resistance to carbenicillin (Conte *et al*, 2015). However, many antibiotics widely used in research have profound effects on mitochondrial function and dynamics, which ultimately affect cell viability, metabolism, and organismal physiology (Houtkooper *et al*, 2013; Moullan *et al*, 2015). Indeed, we observed mildly increased mitochondrial fragmentation in worm tissues exposed to carbenicillin and/or kanamycin (**Fig EV3**). Altogether, these observations suggest that antibiotics might induce mitochondrial fragmentation in *C. elegans*. Therefore, there is a clear need to define cleaner research tools that do not affect the function of mitochondria, as well as greater awareness of the undesired cofounders caused by antibiotics.

Which fluorophore is used can also impact the usefulness of the tool. We observed the development of protein aggregates linked to aging in strains utilizing red fluorophores, irrespective of whether they were tagged to mScarlet, RFP, or mCherry (**Fig EV4**). This occurrence is unique to red fluorophore tags and does not manifest in GFP strains (**Fig EV4**). Moreover, this observations are not contingent on the specific protein to which these fluorophores are attached, as evidenced by the formation of protein aggregates in both the TOMM-20^1-49^::mCherry strain and the TIMM-50::mScarlet (**Fig EV4**). Consequently, in longitudinal studies involving the aging of mitochondrial reporter strains, it is advisable to refrain from using red fluorophores. This phenomenon may provide an explanation for the adverse impacts on growth and reproductive capacity observed in the endogenous TIMM-50::mScarlet strain, effects not seen in TOMM-70 animals with an endogenous GFP tag. Previous research has shown the formation of granular aggregates in immortalized hAECs transfected with mitochondria-targeted mCherry and DSRED (Taiko *et al*, 2022). These aggregates were not due to increased mitochondrial fragmentation or mitophagy, as evidenced by their lack of staining with MitoTracker (Taiko *et al*, 2022). Some cells showed coexistence of both aggregated and properly targeted mCherry and DSRED. This aligns with our findings in *C. elegans*, where the aggregates of fluorophores exhibit increased fluorescence intensity, while the mitochondria show subtle staining.

There is vast potential for advancing tools to investigate organelle biology and inter-organelle communication within mitochondrial research. In this study, we have introduced several novel imaging tools for effectively visualizing mitochondria in *C. elegans*. Nevertheless, we acknowledge the existence of others. For instance, a COX-4::GFP fusion protein strain is commonly used for mitochondrial dynamics studies, where COX-4 is tagged at the C-terminus with eGFP using genome editing methods (Raiders *et al*, 2018). When comparing the fluorescence intensity pattern of the COX-4::GFP strain with our TOMM-70::GFP strain or SKI LODGE TOMM-20 strain, the COX-4::GFP animals exhibit an intensity level intermediate between the other two (**Fig EV5**). In addition to the common use of widely recognized fluorophores like GFP or RFP, there are emerging alternatives. These new fluorophores not only serve the purpose of staining mitochondria but also contribute to evaluating various parameters related to mitochondrial function. This is the case of Supernova, a monomeric variant of KillerRed which allows for the evaluation of ROS levels by chromophore-assisted light inactivation in many living specimens including *C. elegans* (CALI) (Onukwufor *et al*, 2022; Takemoto *et al*, 2013). In addition to this, mt-Keima consists of a pH-dependent fluorescence assay based on the coral-derived protein Keima targeted to the mitochondrial matrix (Sun *et al*, 2015). In this case, mt-Keima allows for the detection of mitophagy by a shift in the excitation wavelength from the physiological pH of the mitochondria (pH 8.0) to the lysosome (pH 4.5) upon engulfment of the mitochondria by the autophagosome (Sun *et al*, 2017, 2015). While not yet implemented in *C. elegans*, this method has demonstrated success in cultured cells (Sun *et al*, 2017) and unfixed mouse tissues (Sun *et al*, 2017, 2015; Katayama *et al*, 2011).

With the relative ease of CRISPR insertion in *C. elegans*, and safe harbor knock-in systems such as SKI LODGE (Silva-García *et al*, 2019), the future is bright for expanding this model system for studying mitochondria, additional organelles, and their interactions. Our comparative analysis highlights the importance of selecting appropriate tools based on specific experimental goals (**Table EV1, Fig EV6**). The significance of these newly generated tools extends beyond this study, as we anticipate their widespread use within the scientific community working on *C. elegans*. We envision that the application of these tools will open avenues for further exploration and discoveries in the realm of mitochondrial research.

## Materials and Methods

### C. elegans strains and husbandry

Worms were grown and maintained on standard nematode growth media (NGM) seeded with *Escherichia coli* (OP50-1), and maintained at 20°C. OP50-1 was cultured overnight in LB at 37°C, after which 100 μL of liquid culture was seeded on plates to grow for 2 days at room temperature. HT115 was cultured overnight in LB containing carbenicillin (100μg/mL) and tetracycline (12.5μg/mL). Then, 100 μL of liquid culture was seeded on NGM plates containing 100 μg/mL carbenicillin. To make WBM1231 (SKI LODGE TOMM-20::GFP) and WBM1232 (SKI LODGE TOMM-20::mCherry), CRISPR/Cas9 was used to insert *tomm-20aa1-49::GFP* or *tomm-20aa1-49::mCherry*, respectively, into WBM1140 (Silva-García *et al*, 2019). To make WBM1444 (TOMM-70::GFP), CRISPR/Cas9 was used to insert GFP on the carboxyl terminus of tomm-70 at its endogenous locus in the *C. elegans* genome. Similarly, mScarlet was inserted on the C-terminus of the *scpl-4* gene (TIMM-50) at the endogenous *C. elegans* locus to generate WBM1688 (TIMM-50::wrmScarlet). To generate the myo-3∷gfpmt(zcIs14) (SJ4103) strain, which we obtained from CGC, the myo-3∷GFP^mit^ plasmid expressing a mitochondrially localized green fluorescent protein (GFP) (GFPmt) with a cleavable mitochondrial import signal peptide (Labrousse *et al*, 1999; Benedetti *et al*, 2006) was integrated stably according to previous instructions. N2 Bristol (wild type (WT) and COX-4::GFP (JJ2586) were obtained from CGC.

### CRISPR/Cas9 gene editing

All gene edits were performed based on previously described protocols (Paix *et al*, 2015; Silva-García *et al*, 2019). Homology repair templates were amplified by PCR using primers that introduced a minimum of 35 base pairs of homology flanking the site of insertion at both ends. CRISPR injection mixes were generated with the following composition: 2.5 μl tracrRNA (4 μg/μl), 0.6 μl dpy-10 crRNA (2.6 μg/μl), 0.5 μl target gene crRNA (2.6 μg/μl), 0.25 μl dpy-10 ssODN (500 ng/μl), homology repair template (200 ng/μl final in the mix), 0.375 μl Hepes pH 7.4 (200mM), 0.25 μl KCl (1M) and RNase free water to make up the volume to 8 μl. Prior to injection, 2 μl purified Cas9 (12 μg/μl) was added before the solution was centrifuged at 13,000 rpm and then incubated at 37 ºC for 10 min. Mixes were microinjected into the germ line of day 1 adult hermaphrodites as previously described (Silva-García *et al*, 2019; Mello *et al*, 1991). Worms generated using CRISPR were outcrossed at least six times before being used for experiments to remove the co-injection marker phenotype and other off-target edits.

### Lifespans

Lifespans assays were performed at 20°C on 6 cm NGM plates seeded with OP50-1, or seeded with HT115 containing 100 μg/mL carbenicillin. All worms were kept fed for at least three generations after thawing to minimize transgenerational effects of starvation on lifespan (Rechavi *et al*, 2014). Then, worms were synchronized through timed egg lays using gravid adults. When progeny from the egg lay reached Day 1 of adulthood, 120 worms were transferred to fresh plates at 20 worms per plate. When excessive censoring was anticipated, the initial population consisted of 160 worms. For the lifespan assays involving HT115 bacteria, worms underwent prior bleaching and were left for 2 generations before initiating the egg lay. When working with genotypes that were developmentally delayed, bleaches and egg lays were staggered so that all worms reached Day 1 adult stage simultaneously with the wild-type animals. Worms were transferred to freshly seeded plates every day until reaching day 4 in order to separate from progeny, and then every other day until day 10 or 11 of adulthood. Survival was assessed every other day, with worms considered dead when unresponsive to 3 touches on the head and tail. Worms were censored if they crawled up the wall, bagged, or exploded. Additional details and statistical information can be found in the Table 2.

### Generation time assay

Animals were synchronized by performing an egg lay 3 days prior to the experiment. The egg lay for TIMM-50 and SJ4103 animals occurred 3 hours earlier than the other strains due to their developmental delay, while the egg lay for all other strains took place in the evening. During the egg lays, 20 animals were placed on 2-day seeded NGM plates, allowed to lay eggs for 30 minutes, and then removed. Early in the morning 3 days later, 15 L4 animals for each condition were selected and placed onto individual 3.5 cm NG plates, which had been seeded 1 day prior with 50 μL of OP50-1 bacteria. Animals were monitored and scored every hour until the first egg was laid.

### Brood size measurement

Animals were synchronized by egg lay onto NGM plates with OP50-1 bacteria following the instructions in “Generation time assay”. At the L4 stage, 15 animals from each condition were placed individually on 3.5 cm NG plates seeded with OP50-1 bacteria. During the consecutive two days, animals were transferred to freshly seeded 3.5 cm NG plates twice per day, once in the morning and then in the evening. During the third and fourth days of the assay, animals were transferred only once. The plates containing the eggs were kept in the incubator at 20°C for 3 days until the progeny developed into day 1 adults. The number of progeny on each plate was documented, and the daily counts were summed for each parent individually.

### Body size measurement

Worms at day 1 of adulthood were anesthetized on NGM plates without bacteria using 1 mg/mL tetramisole. Once still, worms were imaged on a Zeiss Discovery V8 dissection microscope with an Axiocam camera. All animals were imaged in brightfield at 5x with a constant exposure of 65 ms. Body length was analyzed in ImageJ by drawing a line end-to-end down the midline of the worm and measuring the length of the line.

### Confocal microscopy

Worms were anesthetized in 0.1 mg/ml tetramisole in 1X M9 buffer on empty NGM plates and mounted on thin 2% agarose pads on glass slides with 0.05 mm Polybead microspheres (Polysciences) for immobilization. A No. 1.5 cover glass was gently placed on top of worms and sealed with clear nail polish. Images were performed on a Yokogawa CSU-X1 spinning disk confocal system (Andor Technology, SouthWindsor, CT) with a Nikon Ti-E inverted microscope (Nikon Instruments,Melville, NY), using a Plan-Apochromat 100x/1.45 objective lens. Images were acquired using a Zyla cMOS camera and NIS elements software was used for acquisition parameters, shutters, filter positions and focus control. Images were taken from at least 10 worms per condition, with a minimum of 2 independent experiments performed. For the imaging of the endogenous strains where the fluorophore intensity is dimmer, a binning of 2×2 was used. For staining with TMRE (ThermoFisher Scientific T669), wild-type N2 worms were placed on OP50-1 seeded NGM plates with 10 µM TMRE for 24 hours prior to imaging.

### Statistical analysis

Data were graphed and analyzed in GraphPad Prism

9. For lifespan experiments, survival curves were analyzed using the Log-rank (Mantel–Cox) test. A one-way ANOVA followed by a multiple comparisons test was performed to identify statistically significant differences between groups. For all figures: * indicates p<0.05, ** indicates p <0.01, *** indicates p < 0.001, **** indicates p <0.0001.

## Acknowledgments

Funding support was provided by NIH/NIA R01AG044346, R01AG067106 and R21AG056930. P.Y. was funded by F31AG066458. The graphical abstract and Fig EV6 were created with BioRender. We thank the *Caenorhabditis* Genetic Center for providing strains. We also thank the Mair laboratory members for comments and discussion on the project and manuscript.

## Author Contributions

M.V.-A, P.Y. and W.B.M. conceived the idea for the study. P.Y. and A.L. generated the strains for visualizing mitochondria. M.V.-A and S.R.-S. performed the experiments and analyzed the data. M.V.-A., P.Y. and W.B.M. wrote the manuscript, and all authors contributed to revision and edits.

## Expanded View Figures

**Figure EV1.**
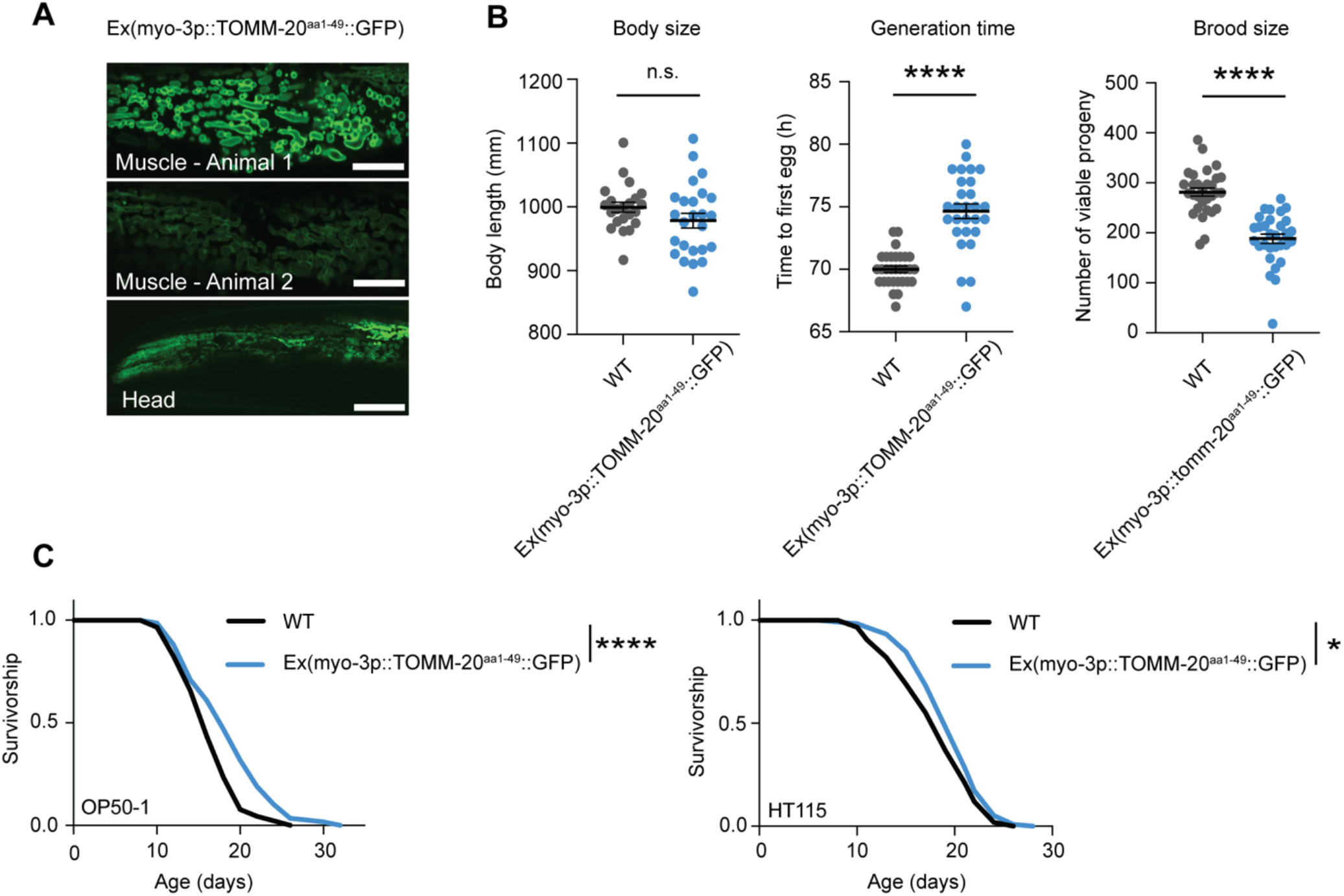
Muscle-specific extrachromosomal array *C. elegans* strain. A. Representative fluorescence pictures depicting mitochondrial networks in the body wall muscle of 2 different worms on day 1 of adulthood. Same laser exposure (300 ms) was used for the three images. Expression of the TOMM-20^aa1-49^::GFP transgene is variable in the progeny from the same hermaphrodite due to the extrachromosomal array. This variability can also be observed in the head section. Scale bar (muscle): 10μm. Scale bar (head): 20μm. B. Healthspan analysis of the extrachromosomal myo-3p::TOMM-20^aa1-49^::GFP strain compared to wild-type N2 worms, showing body size (left graph), generation time (middle), and brood size (right). Body size measurements were performed with 20-25 individual worms. Generation time and brood size measurements were performed with 30 worms pooled from two independent experimental repeats. All values are presented as mean ± SEM. ***p < 0.001 versus the respective WT group (two-tailed Student’s t test) C. Survival curve of the extrachromosomal myo-3p::TOMM-20^aa1-49^::GFP strain compared with WT in OP50-1 (left lifespan) and HT115 (right lifespan) bacteria. *p < 0.05, ****p < 0.0001 indicating significant difference compared to WT animals by one-way ANOVA with Tukey’s multiple comparisons test.

**Figure EV2.**
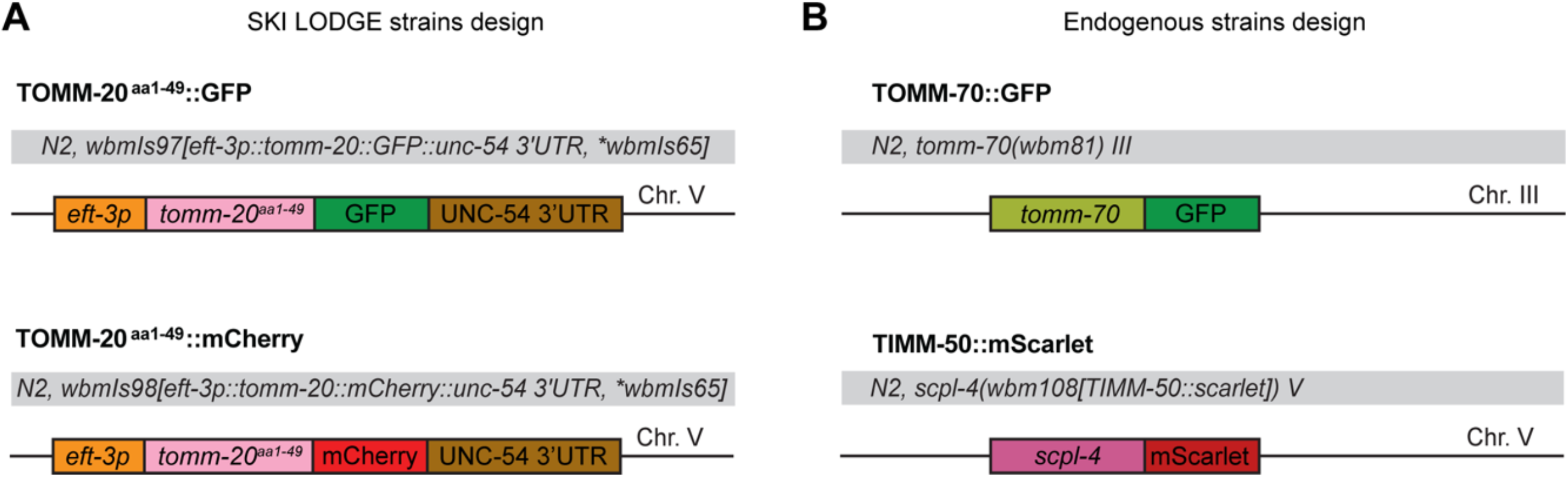
Novel *C. elegans* tool design. A. Schematic of the design for the mitochondrial TOMM-20^aa1-49^ strain with ubiquitous expression (*eft-3* promoter) tagged with GFP or mCherry generated using the SKI LODGE system. B. Schematic of the design for the endogenous strains TOMM-70::GFP and TIMM-50::mScarlet, respectively, using CRISPR/Cas9.

**Figure EV3.**
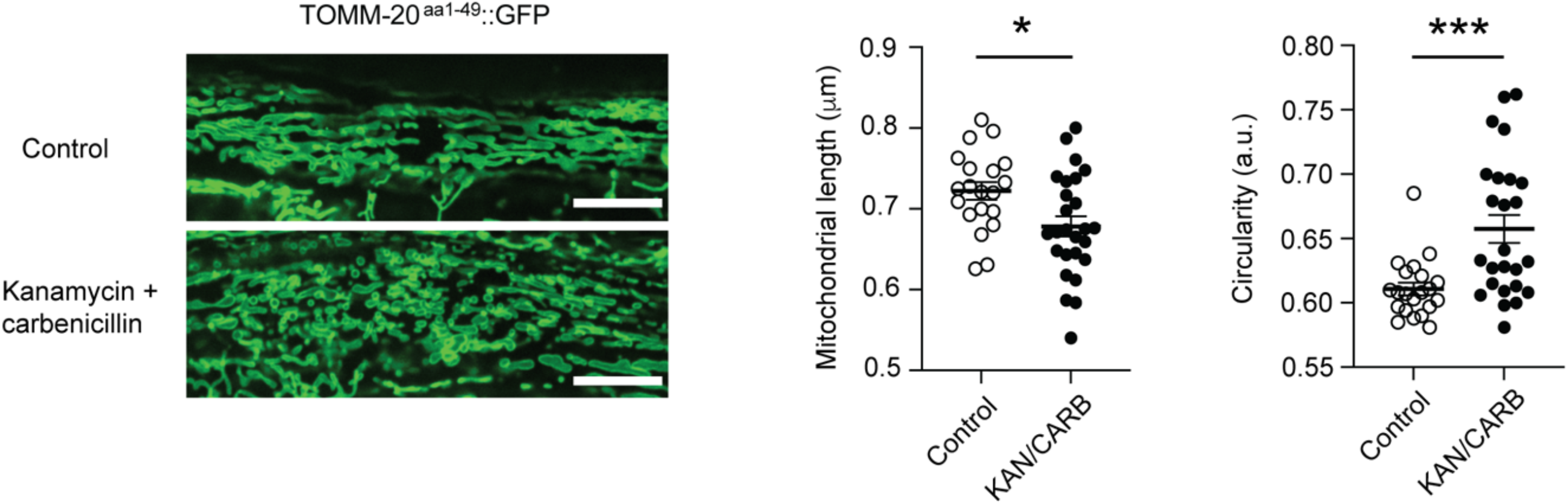
Effect of antibiotics on the SKI LODGE TOMM-20^aa1-49^::GFP *C. elegans* strain. Mitochondria in muscle cells of (SKI LODGE) TOMM-20^1-49^::GFP animals exposed to a combination of the antibiotics kanamycin (0.05 mg/ml) and carbenicillin (0.1 mg/ml) for 24 hours were more fragmented compared to control animals that were not exposed to antibiotics. Representative images from 20-30 animals per condition. Quantification of mitochondrial fragmentation is represented by mitochondrial length and circularity measured by MitoMAPR. *p < 0.05, ***p < 0.001 by two-tailed Student’s t test. Scale bar: 10μm

**Figure EV4.**
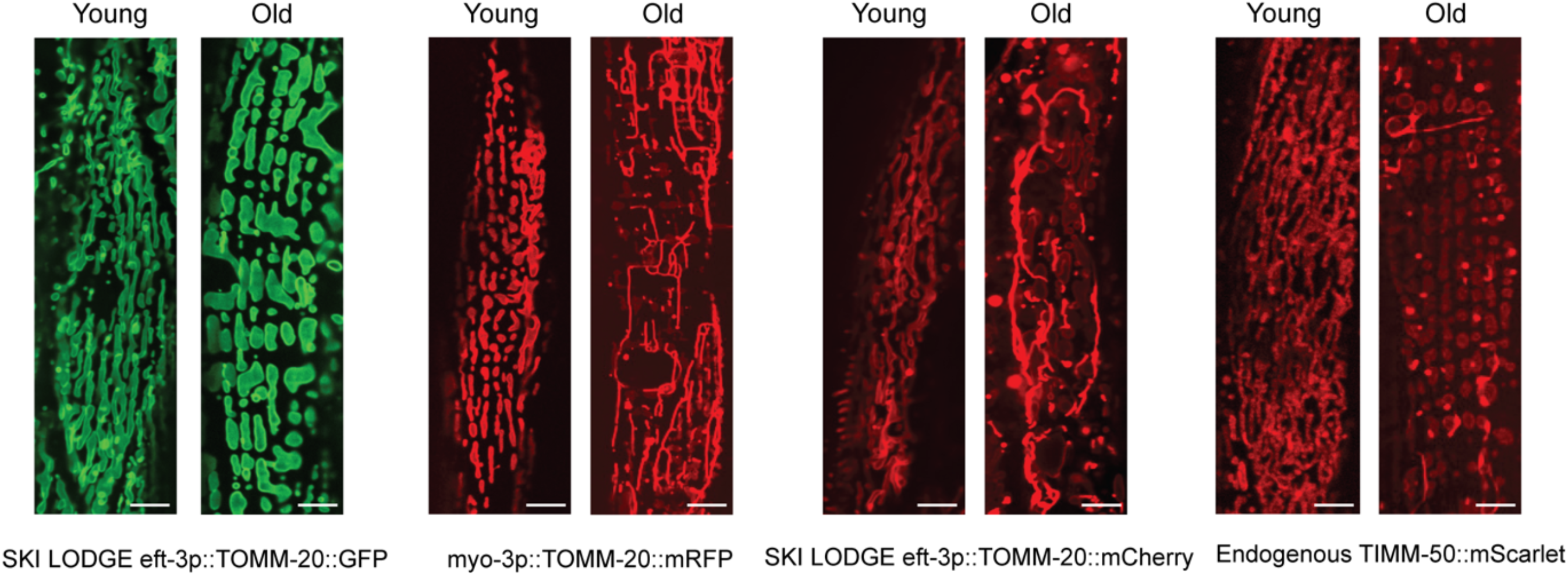
Protein aggregate formation in *C. elegans* strains utilizing red fluorophores. Comparison of young (Day 1) and old (Day 8) worms revealed the development of protein aggregates associated with aging, consistently observed in strains utilizing red fluorophores, including mScarlet, RFP, or mCherry. This phenomenon was exclusive to red fluorophore tags, as TOMM-20^1-49^::GFP animals did not exhibit protein aggregation. Scale bar: 10μm

**Figure EV5.**
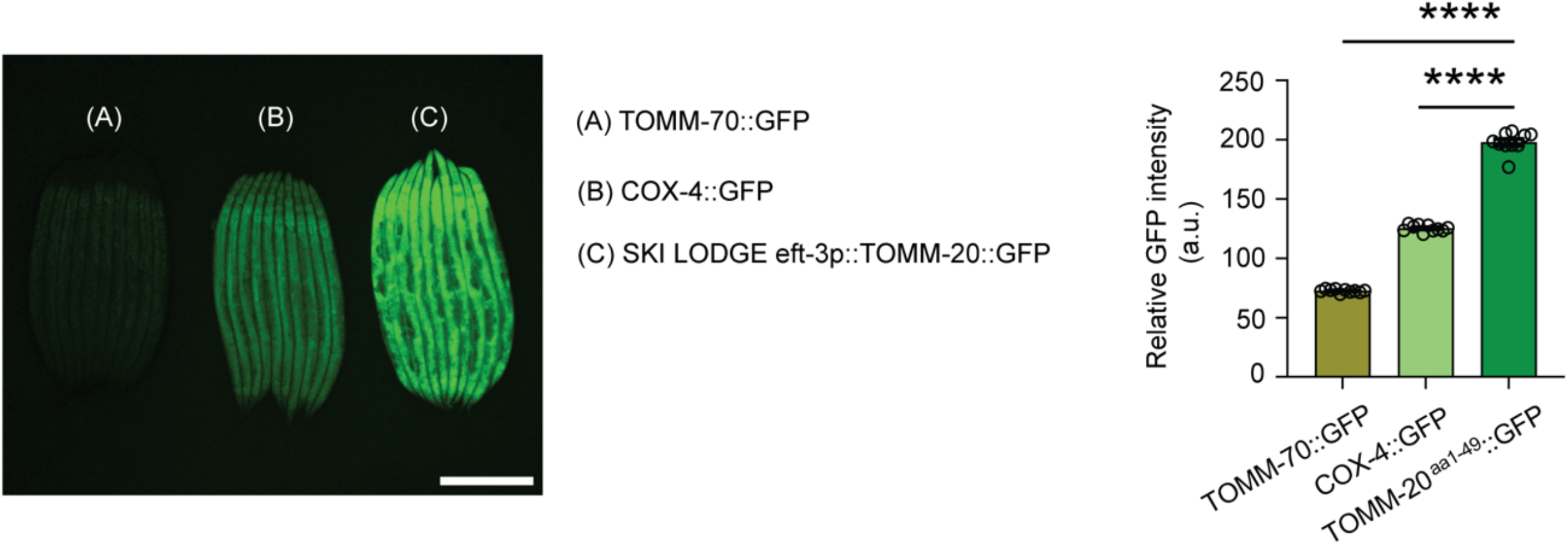
Relative fluorophore intensity in COX-4::GFP *C. elegans* animals. Comparison of the GFP fluorescent intensity of the endogenous TOMM-70::GFP (A), COX-4::GFP (B) and the SKI LODGE TOMM-20^1-49^::GFP (C) animals at day 2 of adulthood. Quantification of GFP intensity levels determines significant difference between the SKI LODGE worms and the endogenous strains. COX-4::GFP animals display intermediate GFP fluorescent intensity levels as compared to the other two strains. One-way ANOVA followed by a multiple comparisons test with the mean of the TOMM-20^1-49^::GFP group was performed to identify statistically significant data. ****p < 0.0001. Scale bar: 500μm.

**Figure EV6.**
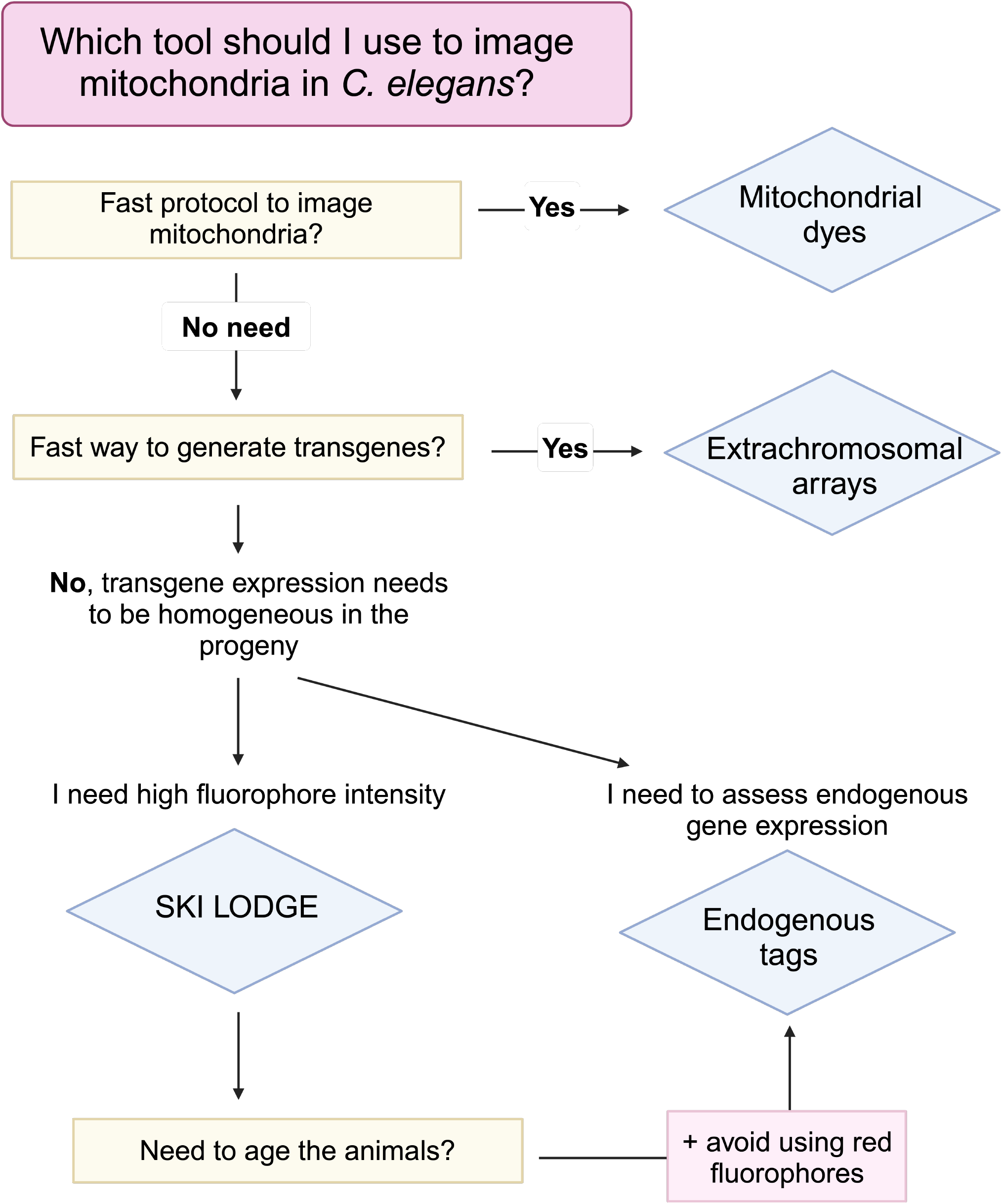
Decision flowchart for imaging mitochondria in *C. elegans*. Schematic guide for selecting the most suitable tools to image mitochondria in *C. elegans* based on experimental goals.

**Table EV1.** Summary of mitochondrial imaging tools in *C*.*elegans* presented in this work.

| Tool | Advantages | Disadvantages |
| --- | --- | --- |
| Mito-GFP (2006) | Established tool | Variable baseline expression levels |
|  | Muscle specific promoter | Animals display developmental delay and reduced brood size |
|  | Widely used | Relies on intact mitochondrial import function for visualization |
| Extrachromosomal Arrays | Reliable means of expressing transgenes | Semistable, requiring selection of transformed animals |
|  | Tissue specific options | Variations in expression level and pattern due to array characteristics |
|  |  | Laborious selection process for transformed animals |
| Mitochondrial dyes (i.e. TMRE) | Convenient means to visualize mitochondria in worm tissues | Staining variability across tissues due to infiltration |
|  | Brief experimental procedures | Off target effects, dye aggregation, or quenching may interfere |
|  |  | Dependency on intact mitochondrial membrane for consistent staining |
| SKI LODGE Strains | Increased fluorophore intensity and homogeneity across tissues | Limited assessment of relative mitochondrial expression levels |
|  | CRISPR assisted integration for stability | Minor changes in lifespan and/or health span indicators |
| Endogenous Strains | Provides insights into endogenous mitochondrial expression levels | Lower fluorophore intensity compared to SKI LODGE strains |
|  | No striking alterations in animal physiology |  |

**Table EV2.** List of *C. elegans* strains.

| Genotype | Strain ID | Alternative name | Source |
| --- | --- | --- | --- |
| wild type | N2 bristol |  | CGC |
| zcIs14[myo-3p::GFP(mit)] | SJ4103 | myo-3::GFPmit | CGC |
| wbmEx289[myo-3p::tomm-20 (aa1-49)::GFP::unc54 3'UTR] | WBM671 | Ex(myo-3p::tomm-20aa1-49::GFP) | Mair lab |
| N2, wbmls97[eft-3p::tomm-20::GFP::unc-54 3'UTR, *wbmls65] | WBM1231 | SKI eft-3p::tomm20::GFP | Mair lab |
| N2, wbmls98[eft-3p::tomm-20::mCherry::unc-54 3'UTR, *wbmls65] | WBM1232 | SKI eft-3p::tomm20::mCherry | Mair lab |
| tomm-70(wbm81) III | WBM1444 | TOMM-70::GFP | Mair lab |
| scpl-4(wbm108[TIMM-50::scarlet]) V | WBM1688 | TIMM-50::mScarlet | Mair lab |
| tomm-70(wbm81[TOMM-70::GFP] III; scpl-4(wbm108[TIMM50::scarlet]) V | WBM1689 | TOMM-70::GFP ; TIMM-50::mScarlet | Mair lab |
| cox-4(zu476[cox-4::eGFP::3xFLAG]) I | JJ2586 | COX-4::GFP | CGC |
| N2, wbmls65[eft-3p::3XFLAG::dpy-10 crRNA::unc-54 3'UTR, V:8645000] | WBM1140 | SKIS: eft-3p::dpy-10 crRNA site | Mair lab |
| syIs268 (myo-3p::TOM20::mRFP) | myo-3p::TOM20::mRFP | myo-3p::TOM20::mRFP | Sean P. Curran |

**Table EV3.**
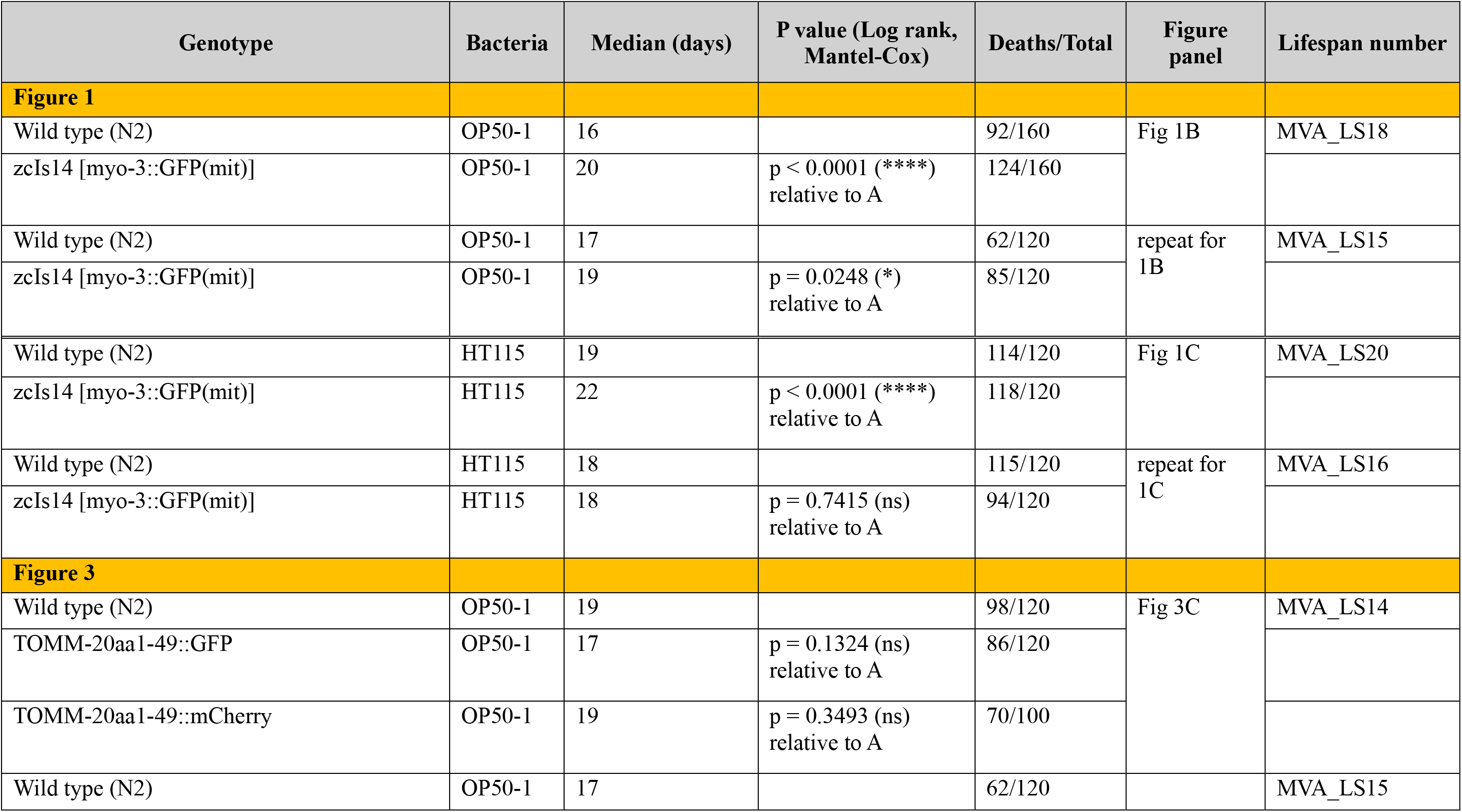
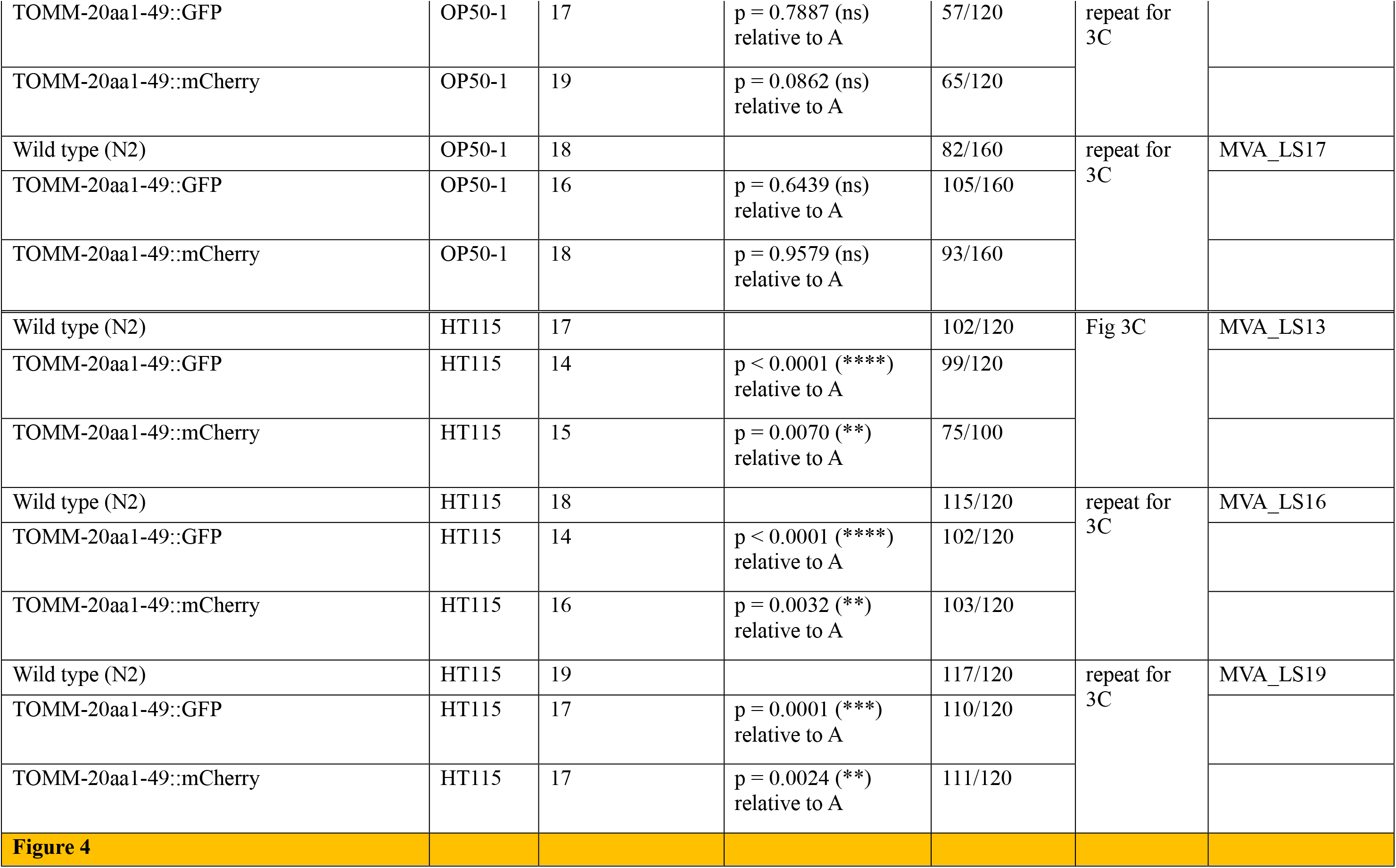
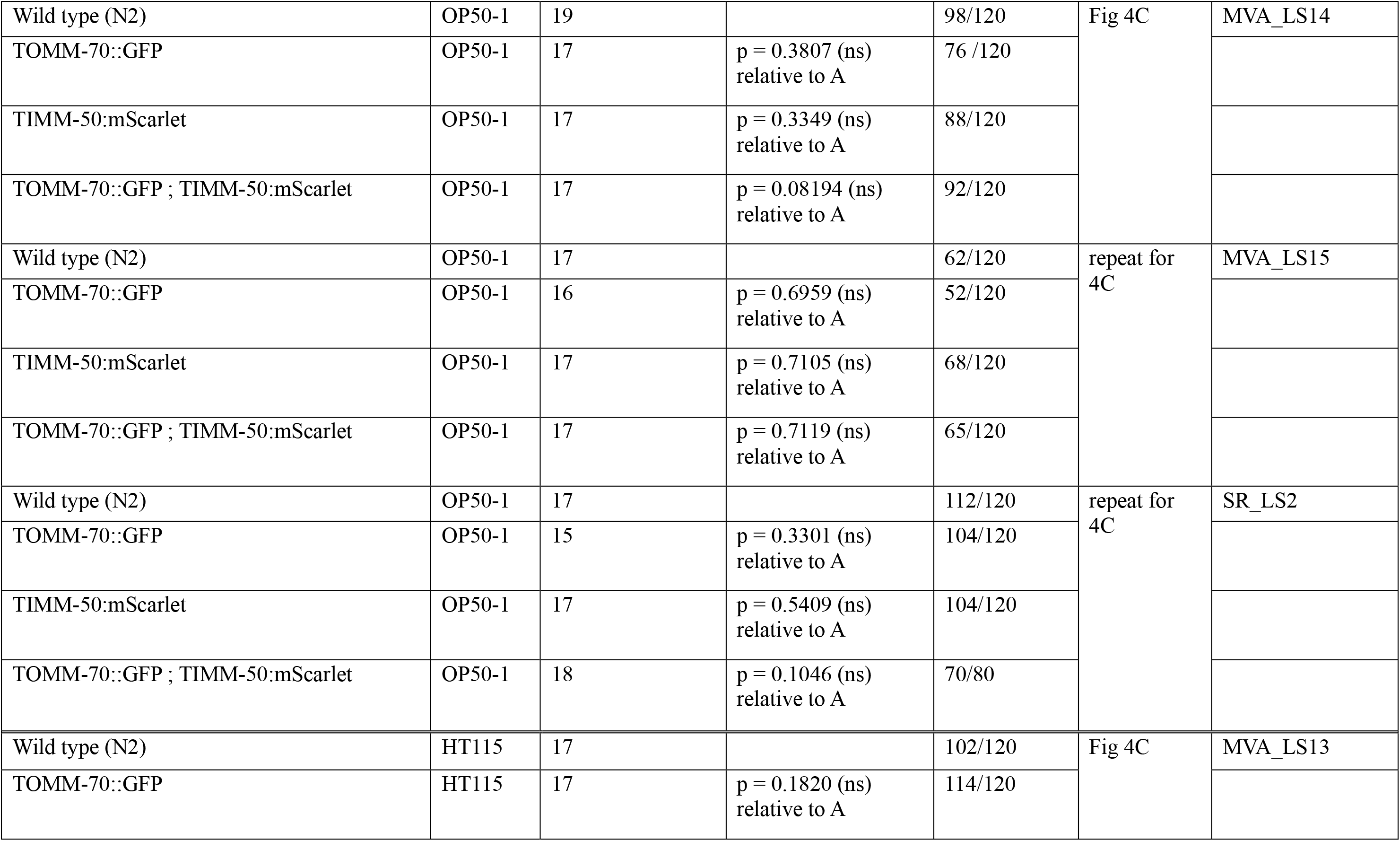
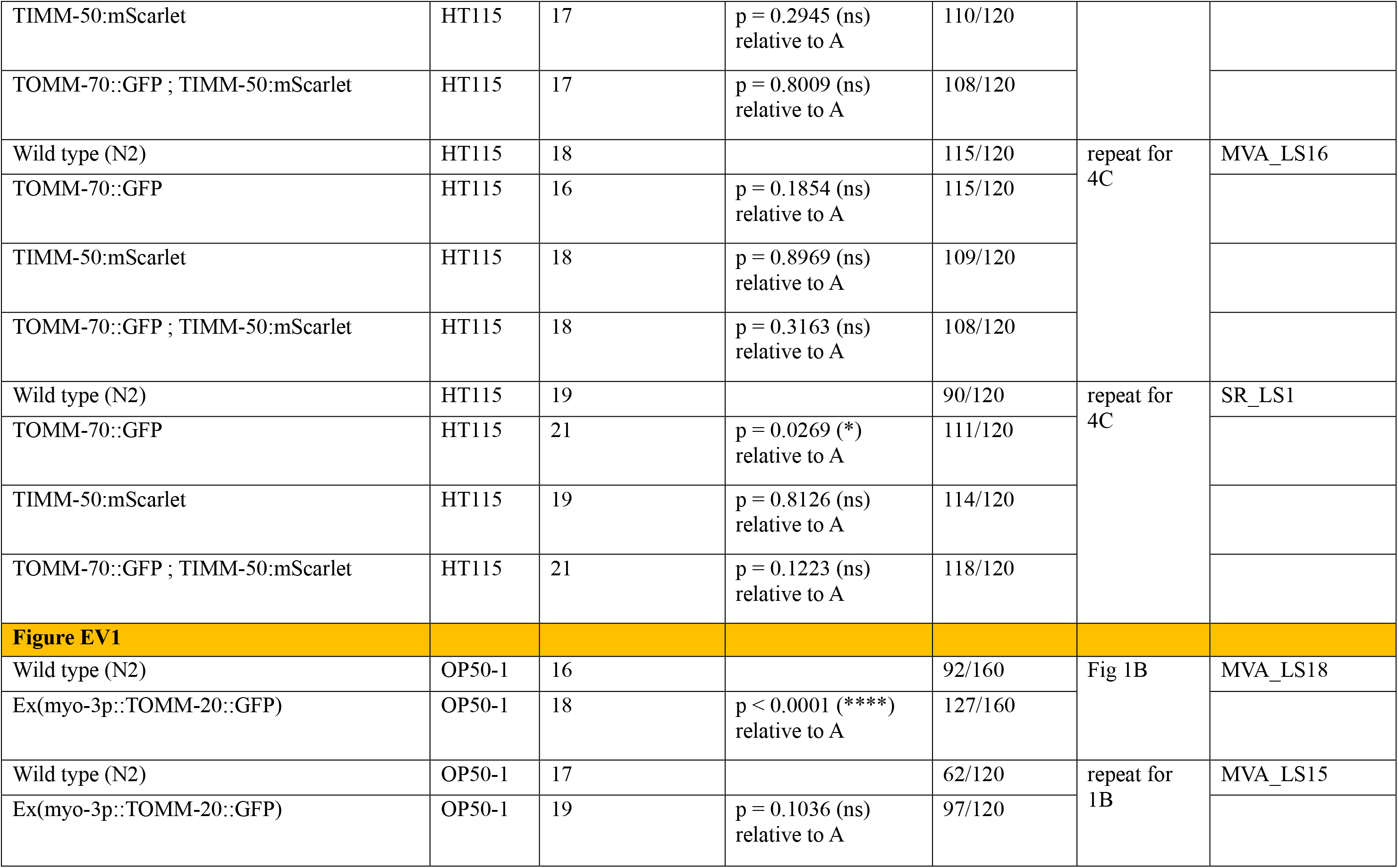
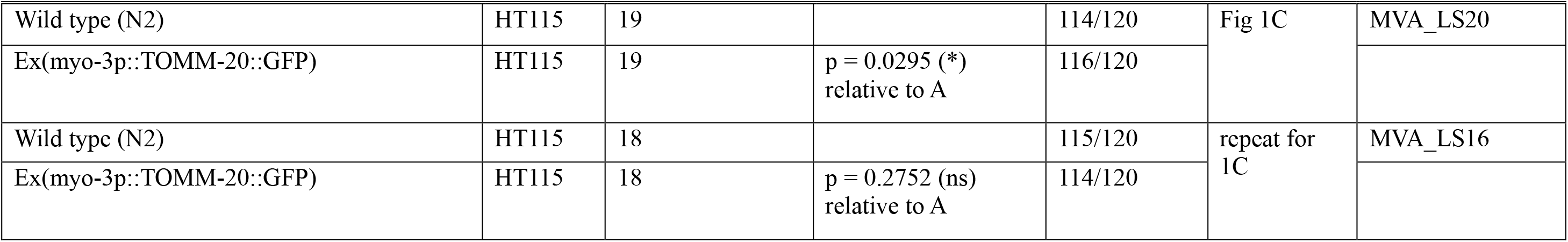
Lifespan table.

